# Analysis of predation-driven inoculum loss and carbon flow in bioaugmented soils through DNA-SIP

**DOI:** 10.1101/2024.04.02.587735

**Authors:** Esteban E. Nieto, Stephanie D. Jurburg, Nicole Steinbach, Sabrina Festa, Irma S. Morelli, Bibiana M. Coppotelli, Antonis Chatzinotas

## Abstract

Bioaugmentation is considered as a sustainable and cost-effective methodology to recover contaminated environments, but its outcome is highly variable. Predation is a key top-down control mechanism affecting inoculum establishment, however its effects on this process have received little attention. This study focused on the impact of trophic interactions on bioaugmentation success in two soils with different pollution exposure histories We inoculated a 13C-labelled pollutant-degrading consortium in these soils and tracked the fate of the labelled biomass through stable isotope probing (SIP) of DNA. We identified active bacterial and eukaryotic inoculum-biomass consumers through amplicon sequencing of 16S rRNA and 18S rRNA genes coupled to modified enrichment factor calculation. Inoculation effectively increased PAH removal in short-term polluted soils but not in long-term polluted soils. A decrease in the relative abundance of the inoculated genera was observed already on day 15 in the long-term polluted soil, while growth of these genera was observed in the short-term polluted soil, indicating establishment of the inoculum. In both soils, eukaryotic genera dominated as early incorporators of 13C-labelled biomass, while bacteria incorporated the labelled biomass at the end of the incubation period, probably through cross-feeding. We also found different successional patterns between the two soils. In the short-term polluted soil, Cercozoa and Fungi genera predominated as early incorporators, whereas Ciliophora, Ochrophyta and Amoebozoa were the predominant genera in the long-term polluted soil. Our results showed differences in the inoculum establishment and predator community behaviours, affecting bioaugmentation efficiency. This highlights the need to further study predation effects on inoculum survival to increase the applicability of inoculation-based technologies.

## Introduction

Pollution is a major consequence of human activities, affecting ecosystem productivity and biodiversity. Bioaugmentation, the introduction of pollutant-degrading microorganisms, is a sustainable and cost-effective approach to improve the pollutant removal capacity of a contaminated matrix [1], particularly of persistent pollutants like polycyclic aromatic hydrocarbons (PAH). The inoculant’s survival and pollutant removal efficiency are closely related to the ability of the inoculant to overcome abiotic (e.g., resource availability) and biotic (e.g., competition) pressures after introduction [2] into the new environment. To date, most studies of these biotic pressures have focused on resource competition and antagonism with the native community as the main barrier to inoculant survival and establishment [3]. While predation of bacteria by larger soil organisms is a documented and common phenomenon [4, 5], the role of predation in inoculant survival has received little attention.

Predation exerts top-down control on bacterial communities, regulating bacterial densities, and potentially affecting the survival of introduced microbes. Predatory protists are often assumed to be the main consumers of bacterial biomass [6, 7]. Protists play a key role in biogeochemical cycles: as their C:N ratios are higher than those of their bacterial prey [8], predation releases nutrients that can be used by other members of the food web [9]. Several bacterial groups also exhibit predatory activity, such as obligate predators of the orders Bdellovibrionales and Vampirovibrionales or facultative predators of the order Myxocococcales and the genus *Lysobacter*, among others. Although there is less data available on predatory bacteria, recent studies have shown that these groups are ubiquitous, and their impact on bacterial communities may be underestimated [7, 10, 11].

The composition of the predator community influences bacterial community assembly and diversity by affecting predator feeding selectivity and prey range [6, 12–14]. Simultaneously, the densities of predatory protists and bacteria closely follow the abundance of their bacterial prey [4, 10, 11, 15]. The inoculation of bacteria may trigger a response of the predator community that could hinder the survival of the inoculant [16], and impact the native community. Bacterial amendments may, however, also result in a bottom-up modulation of protistan communities by affecting protist community structure [17]. Trophic interactions are rarely considered in studies of the fate of microbial inoculants in soil [3], especially in contaminated environments [18] where contaminants also affect the abundance of the native microbial community. Protists are more sensitive to PAH toxicity than bacteria, and their sensitivity varies between the different groups [4, 19, 20]. Community shifts (e.g., resulting from disturbance) can increase invasion success of an inoculum due to the increased prey release from protist predation stress [21], however this interaction is modulated by the strength and the duration of the disturbances [22]. Long-term polluted environments may affect the composition of predator communities by selecting for pollutant-resistant species [23, 24], which in turn may affect the bacterial community’s response to inoculation, and the success of establishment of the inoculum. A few studies have shown a response of the predatory community after a bioaugmentation process [25–27]. These studies demonstrated that predation can lead to a reduction of the inoculum below the detection limit [25], while inhibition of protist activity could increase its survival [26]. Importantly, to date, no studies have identified the predatory groups that interact with the inoculant.

Stable isotope probing (SIP) of DNA can assess trophic linkages between microbes or identify their contributions to specific functions in the environment [28]. SIP has been widely applied to trace carbon transfer in soil food webs [29–32], and to identify microbial degraders in bioremediation processes [33– 38]. Most commonly, a labelled substrate is added to the soil to follow its fate as it is consumed by the soil biota. In contrast, inoculating labelled bacteria into the soil can yield insights into the fate of the inoculant, in particular the identification of predatory bacteria and protists potentially involved in the removal of the inoculant [39–41].

To study the impact of predation on inoculum survival and to identify the predators potentially grazing upon the introduced strains during bioaugmentation, we inoculated a ^13^C-labelled co-culture of PAH-degrading strains *Sphingobium sp. AM* and *Burkholderia sp*. Bk [42], in two soils with different PAH exposure histories (long-term and short-term pollution). We hypothesised that (1) in soils exposed to long-term contamination, predators would respond quickly, limiting inoculum survival and biodegradation efficiency to a greater extent than in the soil exposed to short-term pollution, and that (2) predatory protists and bacteria would benefit via direct grazing on the inoculated bacteria or via carbon transfer in the days following inoculation.

## Methods

### Soil sampling

Two soil types with different contamination histories were selected for this experiment. Soils were characterised using Bouyucus, Walkley – Black, Bray Kurtz and Microkjeldahl methods for textural classification, organic carbon, available phosphorus and total nitrogen respectively. Non-contaminated soil (ST) was collected from an urban park near La Plata city, Argentina (34°51’24.6”S;58°06’54.2’’W) and was a clay loam soil with a pH of 5.8-5.9, 3.60% organic carbon, 6.21% soil organic matter, 0.296% total nitrogen, and 0.00042% available phosphorus. No hydrocarbons were detected. To create soil with a short-term contamination history, ST soil samples were artificially contaminated. The contaminant spiking solution contained 150 mg.kg dry soil^-1^ of fluorene (FLU), 600 mg.kg dry soil-1 phenanthrene (PHE), 100 mg.kg dry soil^-1^ anthracene (ANT) and 150 mg.kg dry soil^-1^ pyrene (PYR) to final concentration of 1000 mg.kg dry soil^-1^. This spiking solution was selected because it contained pollutants that were shown to be degradable by the added inoculum [42] and were also found in the long-term contaminated soil.

A long-term contaminated soil (LT) was collected from a petrochemical plant in Ensenada, Argentina (34° 53′ 19″ S 57° 55′ 38″W). This soil was previously treated by landfarming, with several applications of petrochemical sludge over a 2-year period. Soils were sampled ∼10 years after the cessation of petrochemical sludge treatments, showing a total PAH concentration of 573 ± 138 mg kg-1. LT soil was a loam soil with a pH of 7.71, 2.20% organic carbon, 3.78% organic matter, 0.20% of total nitrogen, and 0.00083% available phosphorus.

### Cultivation of ^13^C-labelled bacteria

The co-culture SC AMBk was used as inoculant, made up of *Sphingobium* sp. (AM) and *Burkholderia* sp. (Bk), in a proportion 65:35 of AM:Bk respectively. This co-culture was previously characterised and demonstrated high PAH-degradation efficiency under laboratory conditions [42]. Each strain was grown in liquid mineral medium (LMM) [43] supplemented with 2 g.L ^-1^ of 99% ^13^C_6_-glucose (Sigma-Aldrich, Munich, Germany) as a sole carbon source. In addition, the same strains were grown in LMM supplemented with unlabelled glucose (^12^C-glucose). After 48 hours of incubation (28°C, 150 r.p.m) cells were collected and centrifuged at 6000 r.p.m for 10 min and resuspended in 5 ml of 0.85% NaCl solution. The 16S rRNA gene of both strains was previously sequenced (Festa et al 2013, Macchi et al 2021).

### Microcosm setup for SIP

Soils were sieved through a 2 mm mesh. Soil moisture was assessed by drying, and the final moisture was adjusted to 20% of humidity. Twelve microcosms were constructed with 50 g of sieved soil in 150 ml Erlenmeyer flasks. Half of the flasks contained LT soil, and the other half contained artificially contaminated ST soils. Microcosms were inoculated with 5*10^7^ CFU.g dry soil^-1^ of SC AMBk (determined by OD_580nm_) as droplets to the microcosms. Half of the ST and LT microcosms each received labelled inoculum, and the other half received unlabelled inoculum as controls. Each treatment was carried out in triplicate, incubated over 30 days at 25°C. Soil moisture was adjusted once weekly to maintain the original moisture content. The microcosms were resampled on days 0 (one hour after inoculation), 7, 15 and 30. Three additional non-inoculated microcosms per soil type were used to assess the intrinsic degradative capacity of the native community.

### PAH quantification by GC-FID

To determine PAH concentration, 3 g of soil were sampled from each microcosm at each time point. Samples were lyophilized (L-3, REFICOR) and three consecutive extractions were carried out, using 9 ml of hexane:acetone 1:1 (v/v). In each step, the hydrocarbons were extracted in an ultrasonic bath (Testlab Ul-trasonic TB10TA) at 40 kHz, 400 W for 30 min [44]. The mixture was centrifuged at 3000 rpm for 10 min (Presvac model DCS-16 RV) and the supernatants were collected in brown glass flasks. Then, samples were resuspended in 1 ml of hexane:acetone and filtered (nylon membrane of 0.45-μm pore size). 5 μl of each sample was injected into a Perkin Elmer Clarus 500 gas chromatograph equipped with a flame ionisation detector and a PE-5HT column. The retention times of the different PAH were determined with a mix of the PAH selected for the spiking solution or the Restek 610 mix PAH standard solution, for ST and LT soil microcosms respectively, and quantified using calibration curves through serial dilutions of the standard solutions.

### DNA extraction and gradient fractionation

Total DNA was extracted from 0.5 g of soil from each triplicate mesocosm at 0, 7, 15 and 30 days using the NucleoSpin™ Soil kit (Macherey-Nagel, Germany), and DNA was quantified with Quant-iT BR ds-DNA assay kit with a Qubit fluorometer (Invitrogen, USA).

Fractionation of DNA from ^13^C-labelled treatments and ^12^C-controls was performed according to [28]. Briefly, 2 μg (ST samples) or 0.7 μg (LT samples) of DNA were added to CsCl solution (1.85 g.ml^-1^) and gradient buffer (0.1 M Tris-HCl, 0.1 M KCl, 1 mM EDTA,pH=8.0) to reach a final concentration of 1.72 g.ml^-1^. The mix was loaded into 5.2 ml QuickSeal Polyallomer tubes (Beckman Coulter Pasadena, USA) and isopycnic density ultracentrifugation was carried out at 44,100 rpm for 36 hours at 20°C in an Optima XPN-80 ultracentrifuge (Beckam Coulter) equipped with VTi 62.5 rotor.

After centrifugation, samples were separated into 12 fractions. Each fraction density was inferred through the refraction index determined by an AR200 digital refractometer (Reichert, Seefeld, Germany). Then, DNA samples were precipitated using polyethylene glycol (PEG)/glycogen method and samples were resuspended in 30 μl of buffer TE. DNA final concentration was determined with Quant-iT HS ds-DNA assay kit with a Qubit fluorometer (Invitrogen, USA). For each treatment, SIP DNA density profiles were created. Equal density ranges were selected for the ^13^C-treatment and ^12^C-control profiles, based on the differences in the amount of DNA between labelled and unlabelled treatments. For each replicate, DNA fractions were pooled in two groups, heavy (1.72-1.735 g.ml^-1^) and light (1.70-1.715 g.ml^-1^) (Figure S1). Because samples from day 30 of LT soils showed a low DNA recovery after fractionation (below 35%), these samples were excluded from the analysis.

### Amplicon sequencing analysis

Bacterial and eukaryotic community composition in pooled fractions of each replicate for all microcosms was assessed by sequencing the V3-V4 hypervariable region of 16S rRNA gene and V8-V9 hypervariable region of the 18S rRNA gene, respectively. The gene regions were amplified with 25 cycles of PCR using the primer sets 16S_Illu_V3_F (TCGTCGGCAGCGTCAGATGTGTATAAGAGACAGCCTACGGGNGGCWGCAG) and 16S_Illu_V4_R (GTCTCGTGGGCTCGGAGATGTGTATAAGAGACAGGACTACHVGGGTATCTAATCC) for the 16S rRNA gene, and 18S_ILLU_1422F (TCGTCGGCAGCGTCAGATGTGTATAAGAGACAGATAACAGGTCTGTGATGCCCT) and (GTCTCGTGGGCTCGGAGATGTGTATAAGAGACAGCCTTCYGCAGGTTCACCTAC) for the 18S rRNA gene. PCR products were checked by gel electrophoresis and sequencing was performed using a MiSeq sequencer (Illumina Inc., San Diego, CA) using a 600 bp kit. Due to low concentration and no PCR amplification, three samples were discarded.

Sequencing data processing was performed in R v 4.3.1 (R Core Team, 2023). The 16S rRNA and 18S rRNA gene sequencing reads were filtered, trimmed, dereplicated, chimera-checked, and merged using the *dada2* package (v 1.22.0) using the following parameters: *TruncLen =* 260, 220; *maxEE=* 4; *trimLeft*= 10 for 16S rRNA gene reads; and *TruncLen =* 220, 190; *maxEE=* 5; *trimLeft*= 10 for 18S rRNA gene reads. Reads were assigned with SILVA classifier v.138 for prokaryotes and v.132 for eukaryotes. Further analyses were performed with the *phyloseq* (v. 1.42.0) and *vegan* packages. 16S rRNA sequenced samples had a range of 67-69417 reads per sample and 18S rRNA samples had a range of 74584-169040 reads, and they were standardised to 22159 and 74584 reads per sample, for 16S and 18S rRNA respectively (function *rarefy_even_depth*, with seed =1). Due to the low number of reads, one sample was discarded for 16S analysis after rarefaction.

### Calculation of taxon-specific enrichment factors in labelled rDNA

To identify both bacteria and eukaryotic taxa actively involved in the assimilation of ^13^C from the labelled strains, we developed a modified version of the enrichment factor (EF) index presented by [30]. EFs were calculated for all genera that had a relative abundance greater than 0.5% on average and a prevalence higher than 6% (i.e., present in at least three samples) across all experimental samples. The enrichment factor was calculated as follows:

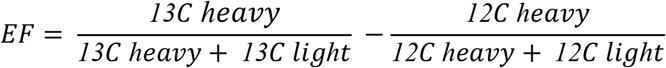

*13C heavy* and *13C light* represent the relative abundance of a genus in the ^13^C-labelled treatment in the heavy and light carbon fractions respectively, and *12C heavy* and *12C light* is the relative abundance of that genus in the two fractions of the ^12^C-controls. Combinatorial subtractions were made between ^13^C treatment triplicates and ^12^C control triplicates, to keep the variability found in each sample. The mean of these values were calculated. The EF ranges between -1 and 1, with EF>0 indicating some degree of enrichment. We set a threshold of EF> 0.015 (i.e., a 1.5% increase in relative abundance of that genus relative to the unlabelled soils) for enrichment and considered those genera that showed significant change of the EF value through time (ANOVA *p*< 0.05).

## Data analysis

Statistical analyses were performed using *rstatix* (v 0.7.2) and PMCMRplus (v 1.9.9). Outliers were identified with the *identify_outliers* function, and Shapiro and Levene tests were done to check for normality and homogeneity of variance. We performed repeated measures ANOVA and post-hoc pairwise comparisons using paired t-tests with a holm correction to determine differences in PAH concentration. To identify changes in relative abundance of *Sphingobium* and *Burkholderia* in the heavy and light fractions of the ^13^C-treatment, we carried out Friedman tests and post-hoc Conover pairwise comparison. Due to the loss of one replicate of the heavy fraction of LT microcosms on day 15, no statistical test was performed. Both % of PAH degradation and relative abundances are reported as *mean ± sd* throughout the study.

In order to distinguish those enriched genera that potential prey on the inoculated genera from those which were enriched by cross-feeding, Kendall correlation analysis was performed between the relative abundance of the inoculated genera and the potential predator, considering a negative value of Kendall correlation coefficient (τ) and statistically significant (p-value < 0.01) for potential predators, while not significant correlation for those enriched by cross-feeding.

## Results

### Bioremediation efficiency

In ST soils, inoculation of the degrading consortium effectively removed all PAHs within the 30 days of the experiment (Figure 1a). Fluorene and anthracene concentrations were below the detection limit after 7 days of incubation, while 98.0 ± 0.7% of phenanthrene (repeated measures ANOVA, *p<* 0.01) and 25.0 ± 12.2% of pyrene (repeated measures ANOVA, *p<* 0.01) were degraded, reaching lower levels than in the non-inoculated microcosms, where no degradation was observed until 15 days of incubation (Figure S2).

**Figure 1.**
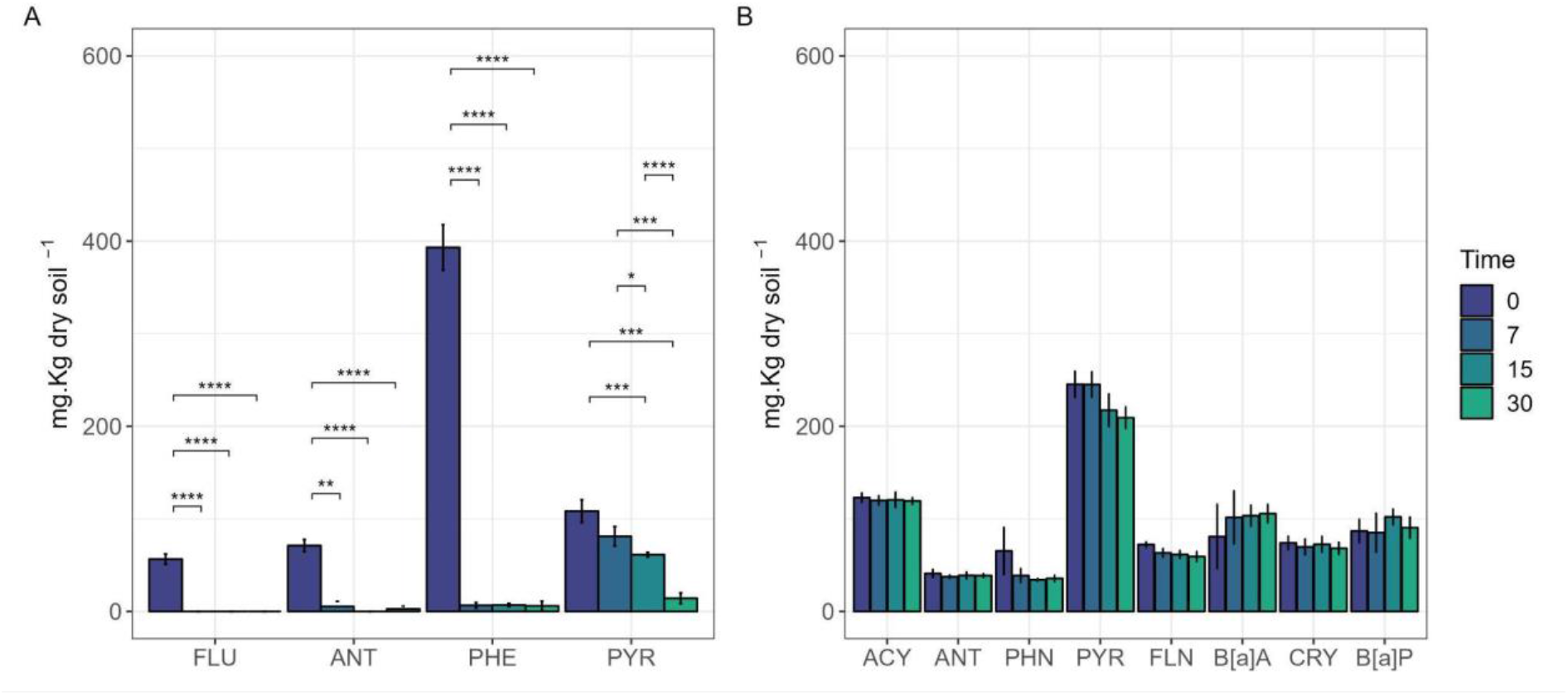
PAH concentrations in ST (a) and LT (b) soil microcosms over incubation period. Results are expressed as the mean concentration of both ^13^C-treatment and ^12^C-control together with their standard deviation.

Eight PAHs were detected in LT soils, including both low molecular weight (acenaphthylene, anthracene and phenanthrene) and high molecular weight PAHs (pyrene, fluoranthene, benzo[a]pyrene, chrysene and benzo[a]pyrene). Notably, inoculating SC AMBk did not reduce the concentration of the PAHs detected in LT soils after 30 days of incubation (Figure 1b, repeated measures ANOVA, *p*> 0.05 for all PAHs)

### Changes in the abundance of the inoculum

We identified two and six ASVs that had a similarity higher than 99.5% with the 16S rRNA sequence of *Sphingobium* sp AM and *Burkholderia* sp Bk, respectively (Table S1 and S2, Figure S3). These ASVs represented, on average, 98.0± 1.8% and 95.3±7.3% of the ASVs belonging to the genera *Sphingobium* and *Burkholderia* in the soils, respectively, and were thus used to estimate the fate of the inoculum. In non-inoculated controls, *Sphingobium* and *Burkholderia* had a relative abundance of < 1% of the total community, on average (Figure S4).

Following inoculation, the initial relative abundance of the genera *Sphingobium* and *Burkholderia* in the heavy fractions of the ^13^C treatment in ST microcosms was 32.2 ± 7.2% and 51.6 ± 6.7% of the community (Figure 2), and gradually dropped (although not significantly; Friedman test, *p=*0.08) to 3.1 ± 0.8% and 8.7 ± 0.5% of the heavy fraction, respectively, on day 30. Notably, the abundance of *Sphingobium* and *Burkholderia* in the light fraction of ST soils increased from 1.7 ± 0.7% and 3.6 ± 1.4% of the light fraction on day 0, to 17.5 ± 1.8 % (Friedman test, p =0.042) and 36.6 ± 0.8% (Friedman test, *p* =0.0293) for *Sphingobium* and *Burkholderia*, respectively, and gradually decreased thereafter.

**Figure 2.**
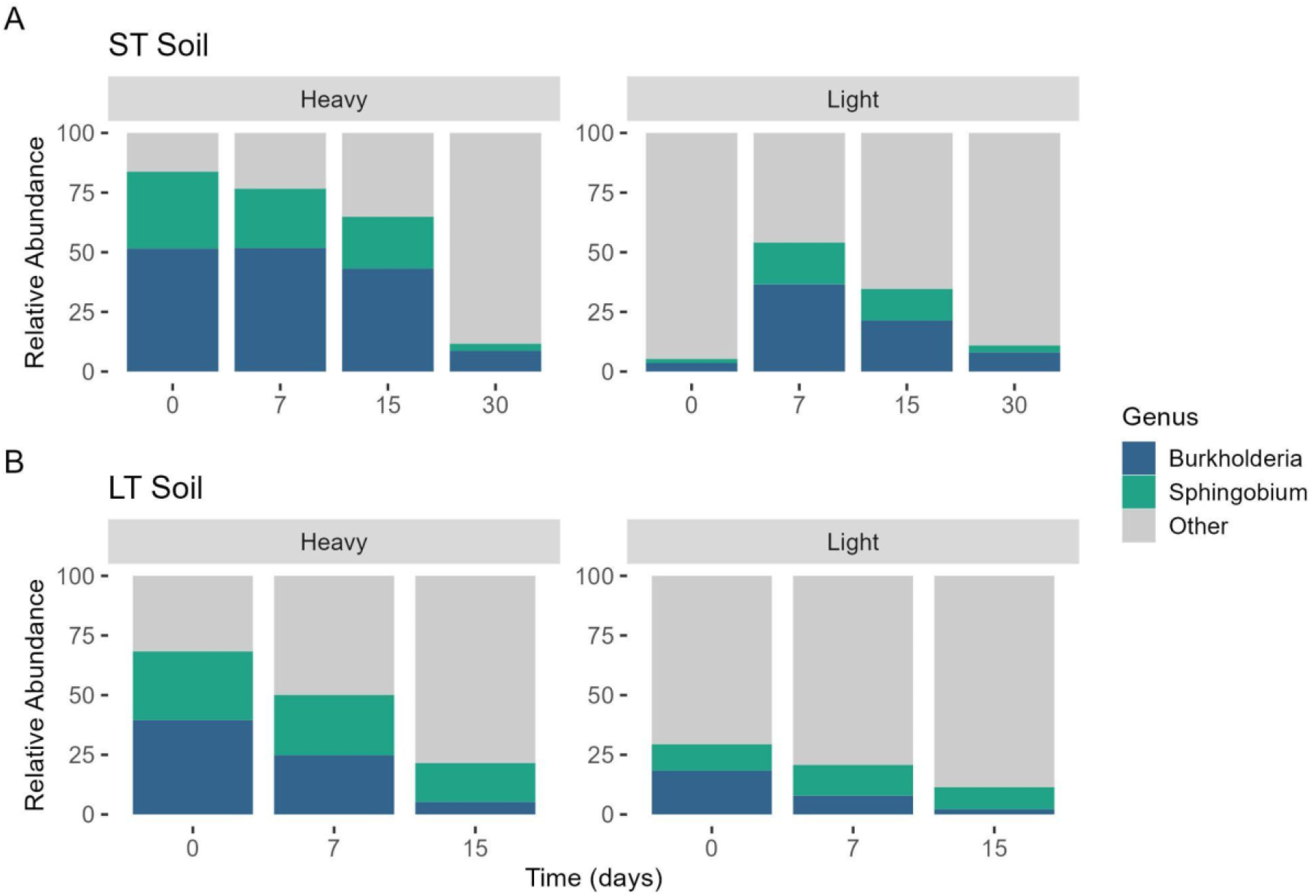
Relative abundance of the genera *Sphingobium* and *Burkholderia* in ST (A) and LT (B) ^13^C treatment soil microcosms over the incubation period. Results are expressed as the mean relative abundance of the biological replicates.

In LT soil, the relative abundance of the inoculated genera in the heavy fraction of ^13^C treatment were 28.8 ± 2.4% and 39.5 ± 2.0% of the total community on day 0, for *Sphingobium* and *Burkholderia*, respectively. However, a decreasing trend was observed through the incubation time, reaching 16.2 ± 0.3% and 5.2 ± 0.1% of the community for *Sphingobium* and *Burkholderia* on day 15. In the light fraction following inoculation, *Sphingobium* and *Burkholderia* made up 10.9 ± 9.5% and 18.4 ± 16.9% of the total community respectively, and no significant changes were detected over time (*Sphingobium:* Friedman test, *p*=0.368; *Burkholderia*: Friedman test, *p*=0.097).

### Enriched community during bioaugmentation

In total, we identified 21 bacterial taxa which were enriched in the ST soils and 3 in the LT soils (Figure 3a). After one hour, *Burkholderia* and *Sphingobium* were the only enriched taxa (EF: 0.42 and 0.46). Their EF decreased during incubation to 0.09 and 0.05 on day 30 respectively. On day 7, *Gemmatimonas* and *Bacillus* were enriched. By day 30, we detected 18 enriched genera. Among them, only *Lysobacter* and a member of Chitinophagaceae family were enriched on day 15, while the remaining genera were enriched on day 30. These genera belonged to Proteobacteria (6 genera) and Acidobacteriota (6 genera), with *Candidatus Udaeobacter* and *Sphingomonas* exhibiting increased relative abundances and the highest enrichment (0.17 and 0.11 respectively). No correlation was found between the relative abundance of the inoculated genera and seven of the enriched genera identified in the ST soil (Table S3). In LT soil, *Promicromonospora* was enriched only after one hour of incubation, while the ASV affiliated to *67-14* (unclassified Solirubrobacterales) and *WN-HWB-116* (unclassified Gammaproteobacteria) were enriched on day 15 and none of these genera correlated with the relative abundance of the inoculated genera (Figure 3b, Table S3).

**Figure 3.**
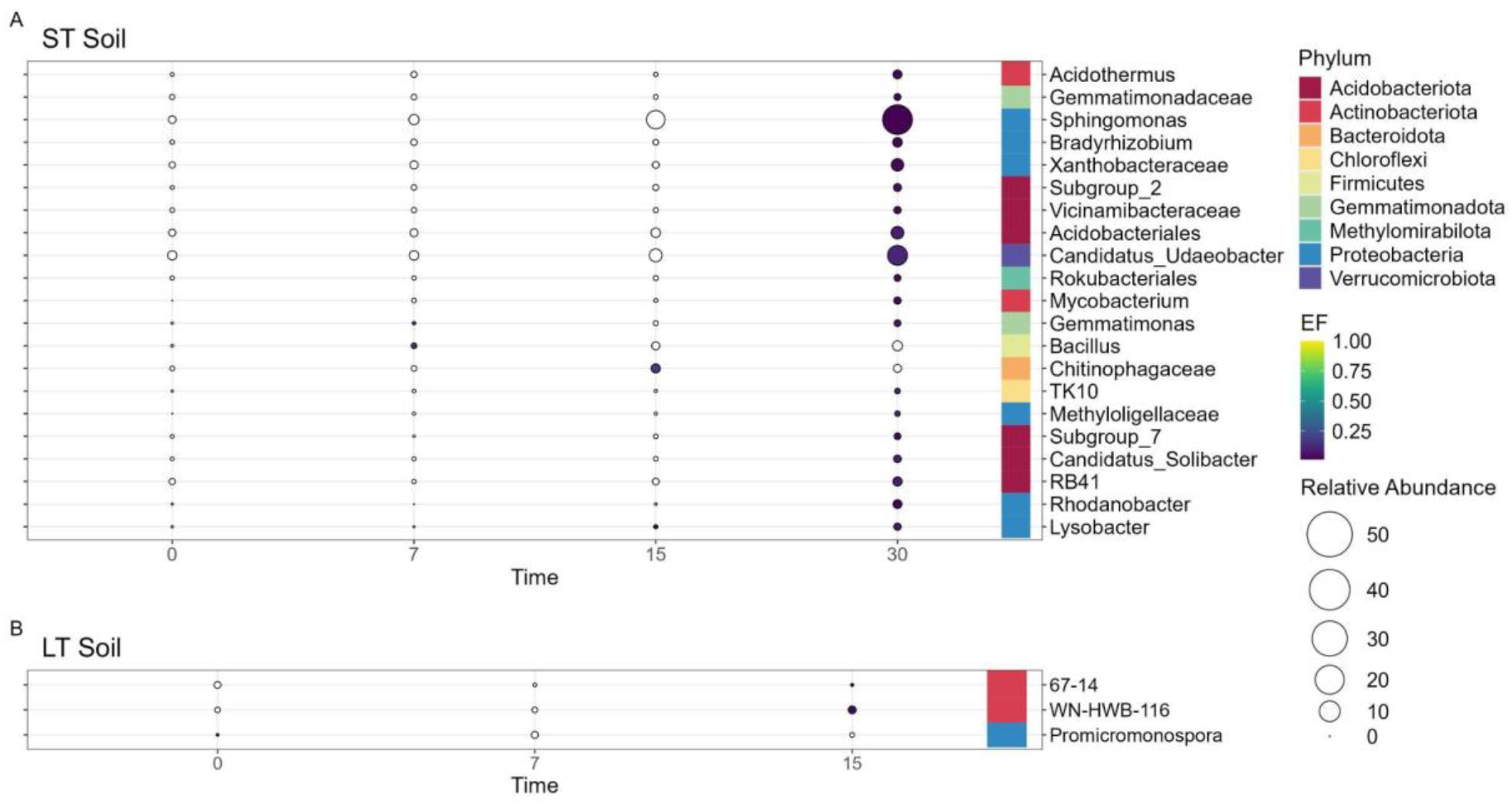
^13^C enriched bacterial genera identified by calculating the EF in the ST(A) and LT(B) soil amended with the labelled consortium SC AMBk after 0 (after one hour of incubation), 7, 15 and 30 days of incubation. The colour scale reflects the mean value of the EFs and the size scale represents the mean relative abundance of each genus in the heavy fraction of the ^13^C treatment. The figure shows the genera whose EF was higher than 0 at least at one time point, showed significant change in their value through time (*p<*0.05), and whose relative abundance was higher than 0.5% in at least one time point. White circles represent EFs lower than 0.01.

We further identified 15 enriched eukaryotic taxa in ST soils (Figure 4a). After one hour of incubation five genera were enriched, including Cercozoa (2 genera), Amoebozoa (1 genus) and Fungi (2 genera). On day 7, eight genera were enriched, which included Cercozoa (4 genera), Amoebozoa (2 genera) and Fungi (2 genera). On day 15, ten enriched genera were identified, among which *Allas* showed the highest EF (0.42) and *Mortierella* and *D3P05A02* (Apicomplexa) showed the highest relative abundance. Only six genera were enriched on day 30, which belong to Cercozoa (4 genera) and Amoebozoa (2 genera). Notably, *Allas* and *Cercomonas* exhibited a negative correlation with the inoculated genera (*Allas:***τ** = -0.71, *Cercomonas:* **τ**= -0,67 for *Burkholderia*, and *Allas:* **τ**=-0.75 for *Sphingobium*; p-value< 0.01, Table S4) Among the eukaryotic enriched genera in LT soil (Figure 4b), *D3P05A02* (Apicomplexa) and *Ophistonecta* were enriched after one hour. On day 7, five genera were enriched, which included Ochrophyta (2), Amoebozoa (1), Ciliophora (1) and Chlorophyta (1). Of these, *Spumella* and *Clamydomyxa* had the highest EF values (0.24 and 0.17 respectively). On day 15, only *Colpoda* remained enriched (EF 0.20). None of the enriched genera showed significant correlation with the inoculated genera (Table S5)

**Figure 4.**
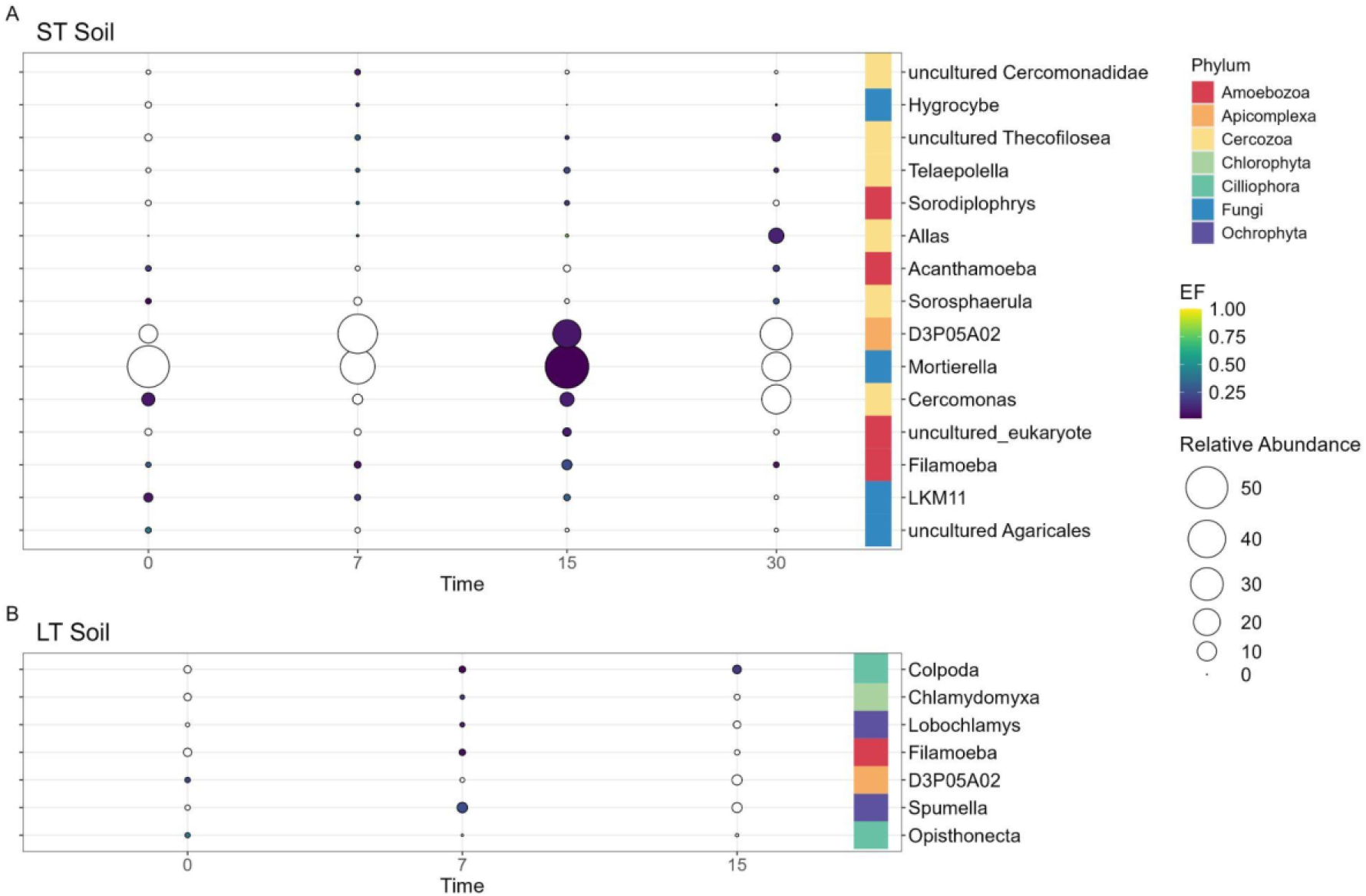
^13^C enriched eukaryotic genera identified by calculating the EF in ST(A) and LT(B) soil amended with the labelled consortium SC AMBk after 0 (after one hour of incubation), 7, 15 and 30 days of incubation. The colour scale reflects the mean value of the EFs and the size scale represents the mean relative abundance of each genus in the heavy fraction of the ^13^C treatment. The figure shows the genera whose EF was higher than 0 at least at one time point, showed significant change in their value through time (*p<*0.05), and whose relative abundance was higher than 0.5%. White circles represent EFs lower than 0.01.

## Discussion

Bioaugmentation is considered a cost-effective and sustainable technology for the removal of pollutants from contaminated environments [1], however the efficacy of this process is highly variable [45–47]. The establishment of inoculated microorganisms is a key part of this process, which may ensure the successful removal of pollutants. Using DNA-SIP to track the incorporation of isotopically labelled biomass carbon from the inoculum strains into members of the soil microbial food web, we demonstrate how the performance of a PAH-degrading two-strain inoculant depends on contamination exposure times, and highlight the role of inoculum predation by the native community in modulating this performance.

Although predation is an important top-down regulation process [4], its impact in bioremediation has been overlooked. To date, few studies consider the role of predation in degradation, and those which do, report both a decreased degradation [25, 26, 48], or increased degradation in the presence of predators [49–51]. Nevertheless, how predator communities vary in different environments, and how this affects the success of bioinoculants remains a significant hurdle to the wider application of this promising technique. These studies lack the identification of the main predatory groups involved in the process.

We found that inoculum biomass was assimilated by members of the native soil microbial communities, and that these differed between soils with different contamination legacies, highlighting the complex relationship between the native microbiota, the pollutant, and the efficacy of bioinoculants in removing it from the environment. We were able to identify the main groups involved in the assimilation of inoculum biomass. In both soils, eukaryotic genera predominated as consumers of the ^13^C-labelled biomass at the first sampling time points, but the identity of individual genera varied between the soils. In the ST soil, members of the Cercozoa phylum and fungal genera were predominant among the enriched genera. This trend continued throughout the month of study, and during this time, additional cercomonadid and amoeboid genera incorporated the labelled biomass. On the other hand, in LT soil, no Cercozoa genus was enriched, and members of Amoebozoa, Ochrophyta and Ciliophora were found among the early incorporators. Previous studies have reported a quick response of the fungal community after the addition of labelled carbon substrates [30] and labelled bacteria [39, 52, 53]. Although saprophytic fungi are considered to mainly decompose plant litter, they can be dominant in the decomposition of newly added residues, including bacterial biomass. Fungi produce a variety of extracellular enzymes attacking compounds not easily degradable (e.g. bacterial cell wall components), releasing low molecular decomposition products available for other organisms [52, 54]. Regarding protists, consumers are the predominant functional group in soil, with Cercozoa being the most abundant group followed by Ciliophora [55]. Due to the rapid division times, they can respond quickly to changes in the prey abundances [4, 15, 56].

Our study suggests that inoculation triggered distinct successional trajectories in the community, between the two soils with different contamination exposure times. This is likely due to the different contaminant tolerance and preference ranges of predatory eukaryotes in the soils with different contaminant legacies, resulting in different communities of potential predators to consume the inoculum. Cercozoa has been documented to dominate soils freshly contaminated with PAH [20]. Also, a positive correlation was found between this group and PAH-degrading bacteria and fungi [57]. However, this study showed that Cercozoa abundance was negatively correlated with high molecular weight -PAH, the main compounds found in the LT soils. Unlike Cercozoa, positive correlations were found between the high molecular weight-PAH concentration and the abundance of Stramenopiles (which includes Ochrophyta) and Amoebozoa [57]. Furthermore, higher abundance of amoeboids in soils chronically contaminated with PAH were previously observed [58, 59]. As with Cercozoa, this pattern is likely related to the higher resistance to PAH toxicity of amoeboids compared to flagellates [19]. However, there is yet a lack of information regarding the interaction between soil protists and pollutants [18], and more studies should be done to confirm these patterns.

In contrast to eukaryotic genera, few bacterial genera were enriched at the first sampling time. The number of enriched bacterial genera increased only by the 30th day, and only in the ST soil. Enrichment of members of the genera *Sphingomonas, Lysobacter, Rhodanobacter*, uncultured Xanthomonadacea has been also shown in previous work, where ^13^C labelled *Pseudomonas putida* and *Achromobacter globiformis* were inoculated in bulk or rhizosphere soils [40]. Also some of the enriched genera (e.g. *Bacillus*) were previously identified as consumers of dead biomass derived from *E. coli* [60]; however, among them, only *Lysobacter* is known as a facultative predator [61]. This fact, together with the late incorporation of labelled biomass, suggests that the enrichment of bacteria may have resulted from cross-feeding [38]. It has been recently suggested that the role of predatory bacteria may be underestimated and that they could play a major role in the soil food web [7, 11]. Interestingly, and contrary to our expectation, we did not identify known predatory bacteria in the first stages after incubation. In both soils, eukaryotic predators were likely the main consumers during the first stages after inoculation, releasing nutrients that could later be used by the bacterial and fungal communities [9].

We also found that *Sphingobium* and *Burkholderia* showed different survival regarding the soil in which they were inoculated, which could also explain the different degradation performances. In the ST soil, both inoculated genera showed a high relative abundance in the heavy fractions of the ^13^C treatments during the first 15 days of incubation. Interestingly, we observed growth in situ for both genera 7 days after inoculation, as indicated from the dilution of the label and the increase of the relative abundance in the light fractions. In contrast, in the LT soil we did not observe any increase in relative abundance in the light fraction, but by day 15, the relative abundance of both genera had progressively decreased in the heavy fraction, meaning the inoculated genera were not able to grow during the incubation period.

Both environmental factors (e.g. resource availability, pollutant toxicity) and antagonistic biological interactions (e.g., competition, predation) can hinder inoculum establishment [2]. Although we did not directly assess competition, in ST soil, the inoculated genera outcompeted the native community in PAH-consumption, as reflected by the increase in degradation, and were able to grow [2], outpacing predation pressure [62]. By the end of the incubation time, PAH concentrations had decreased and were not renewed, but the predation pressure remained, possibly explaining the decrease in the relative abundance of the inoculum at the end of the experiment. In contrast, no degradation was observed in the LT soil. The low survival of the inoculum in LT soil may be the result of higher predator pressures in these soils in addition to the lower PAH-bioavailability [63, 64]. In this scenario, where no growth was observed, *Sphingobium* showed higher survival than *Burkholderia*. Several predation-resistance strategies have been described for *Sphingobium* genus, such as aggregate formation [65, 66] and the presence of sphingolipids which stabilise the outer membrane of the cell wall and reduce digestibility [67, 68].

Combined strategies of bioaugmentation and surfactant-enhanced bioremediation have been proposed as an option to increase the degradation of PAH in environments with low bioavailability [69]. To date, most studies have focused on the degradation efficiency of inocula to design consortia [70–72]. However, it is currently highlighted the need to include other ecological traits [2, 47], particularly predation resistance in inoculum design [3]. Understanding predation interactions between an inoculum and native communities under changing environments will help to increase the probability of successful application of inoculum-based technologies.

## Conclusion

Our work suggests that predation pressure is central to the establishment and success of bioaugmentation in soil, and that predation pressure is also modulated by the environment. At the same time, we find commonalities across soils, including a quick response of the eukaryote-dominated predator community. Although we did not monitor the absolute abundance of the eukaryotic predators, it highlights the relationship between growth and predation, and its impact on inoculum survival. Future research should focus on understanding predation preferences and selectivity by soil microbial eukaryotes in situ and in isolation.

## Declarations

### Ethics approval and consent to participate

Not applicable

### Consent for publication

Not applicable

## Availability of data and material

16S rRNA and 18S rRNA gene sequencing raw data were deposited into NCBI Sequence Read Archive (SRA) under the accession number PRJNA1089553

## Authors’ contributions

EEN, AC and BMC were responsible for conceptualization. EEN performed microcosm set up and sampling. EEN and NS performed DNA ultracentrifugation, fractionation and precipitation. Library preparation was performed by NS. SDJ and EEN performed bioinformatics analyses, enrichment factor conceptualization, and statistical analyses. EEN, SDJ and AC wrote the manuscript, and all authors contributed to its improvement. All authors approved the submitted version

