## Supplemental material for "Analysis of predation-driven inoculum loss and carbon flow in bioaugmented soils through DNA-SIP"

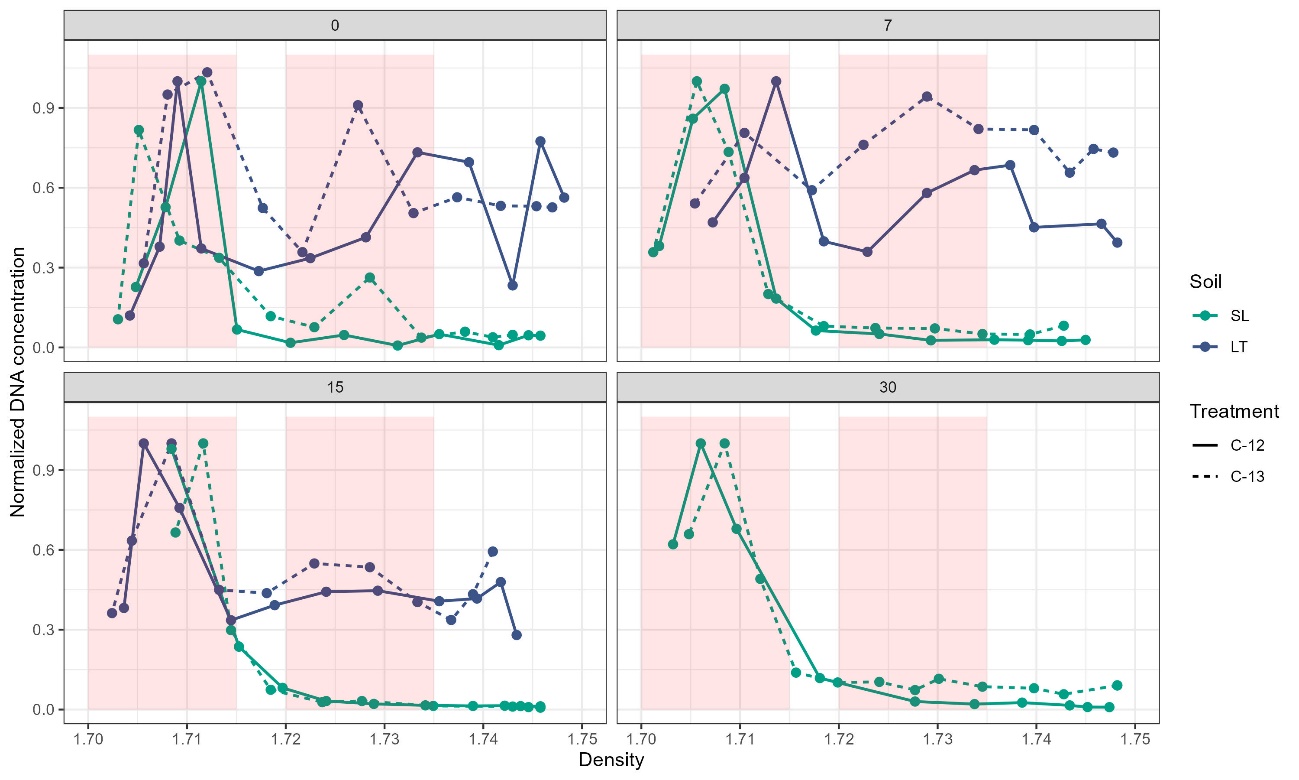

Figure S1: Normalized DNA concentration according to the density of each fraction for both ST (*green lines*) and LT (*blue lines*). Each point represents the mean concentration of the triplicates of ^c13^treatment (*dashed lines*) and ^c12^control (*continuous lines).* Red areas indicate density ranges of heavy and light pools.

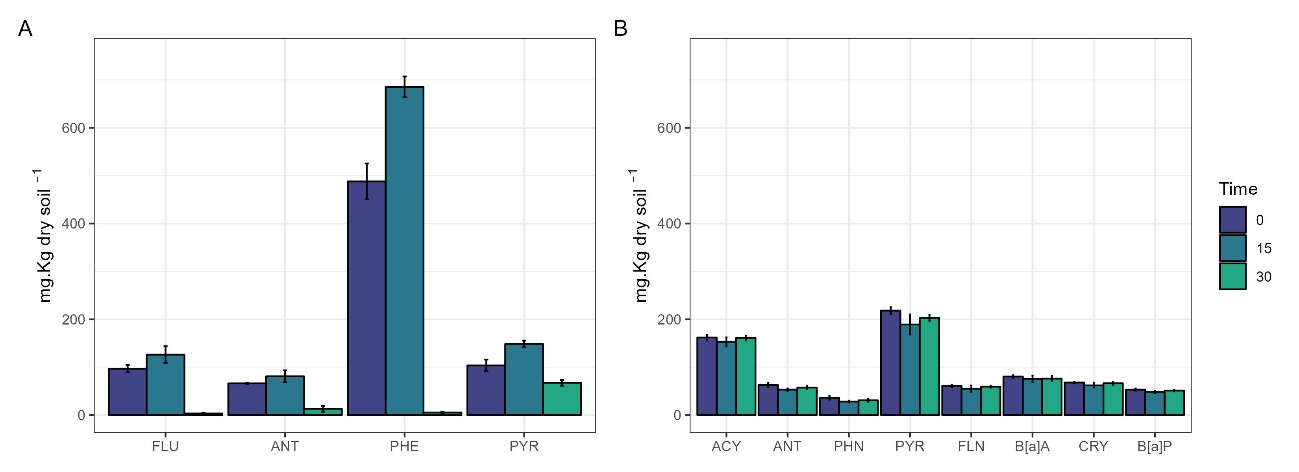

Figure S2: PAH concentrations in ST (a) and LT (b) non inoculated soil microcosms through incubation period. Results are expressed as the mean concentration with their standard deviation.

| ASV | % identity |
| --- | --- |
| ASV4  ASV3 | 99.762  99.524 |

Table S1: Predominant ASV (relative abundance > 0.05%) with the highest percentage of identity with *Sphingobium* sp. AM 16S rRNA gene sequence.

| ASV | % identity |
| --- | --- |
| ASV2  ASV76  ASV70  ASV1  ASV69  ASV58 | 100.000  99.775  99.775  99.775  99.551  99.551 |

Table S2: Predominant ASV (relative abundance > 0.05%) with the highest percentage of identity with *Burkholderia* sp. Bk 16S rRNA gene sequence

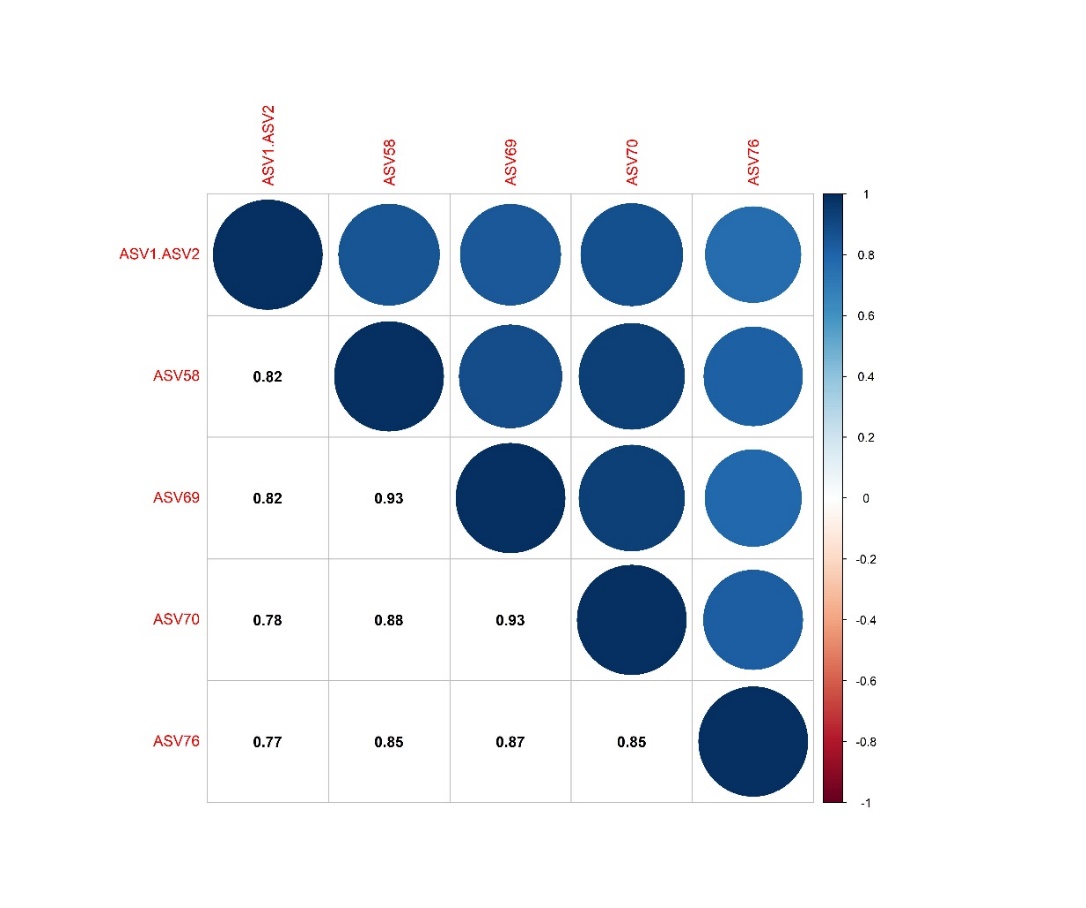

Figure S3: Correlogram of the ASV identified for *Burkholderia* Bk strain using Spearman correlation method.

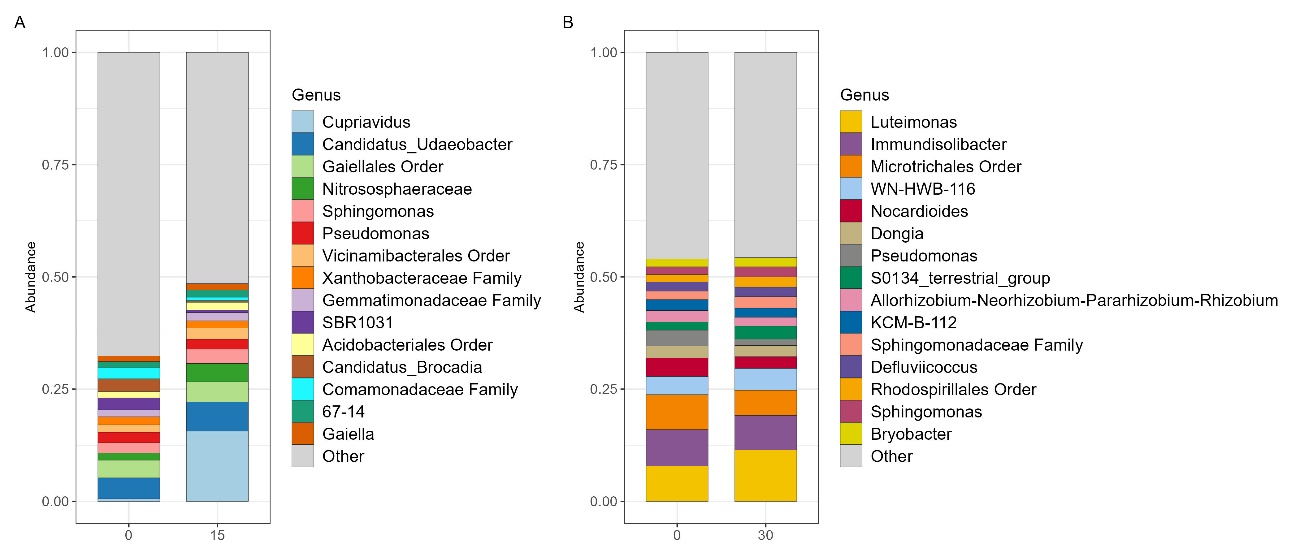

Figure S4: Relative abundance of the top 15 bacterial genera in ST and LT non inoculated soil microcosms. Results are expressed as the mean relative abundance of triplicates at time 0 and 15 days or 30 days of incubation for ST and LT soils respectively.

|  | Correlation with Burkolderia abundance | Correlation with Sphingobium abundace |
| --- | --- | --- |
| Sphingomonas | -0,76*** | -0,82*** |
| Bradyrhizobium | -0,64*** | -0,82*** |
| Acidobacteriales | -0,82*** | -0,82*** |
| Rokubacteriales | -0,58** | -0,82*** |
| Subbroup_2 | -0,79*** | -0,79*** |
| Gemmatimonas | -0,73** | -0,79*** |
| Candidatus_Udaeobacter | -0,88*** | -0,76*** |
| Lysobacter | -0,79*** | -0,73** |
| Mycobacterium | -0,57** | -0,73* |
| Xanthobacteraceae | -0,48* | -0,73*** |
| Vicinamibacteraceae | -0,67** | -0,73*** |
| Candidatus_Solibacter | -0,60** | -0,72** |
| Methyloligellaceae | -0,67** | -0,70** |
| RB41 | -0,79** | -0,67** |
| Gemmatimonadaceae | -0,52* | -0,64** |
| Bacillus | -0,64** | -0,64** |
| Subgroup_7 | -0,69** | -0,63** |
| Rhodanobacter | -0,53* | -0,59* |
| Acidothermus | -0,42 | -0,55* |
| TK10 | -0,51* | -0,54* |
| WN-HWB-116 | -0.42 | -0.50 |
| 67-14 | -0.57 | -0.50 |
| Promicromonospora | -0.50 | -0.48 |
| Chitinophagaceae | -0,36 | -0,48* |

Table S3: Kendall correlation coefficient for the enriched bacterial genera abundance regarding Burkholderia and Sphingobium abundances in both ST and LT soils. **p <* 0.05, ***p <* 0.01, ****p <* 0.001.

|  | Correlation with Burkolderia abundance | | Correlation with Sphingobium abundace |
| --- | --- | --- | --- |
| Allas | | -0,71** | -0,75** |
| Cercomonas | | -0,67** | -0,38 |
| Acanthamoeba | | -0,56 | -0,35 |
| Sorodiplophrys | | -0,31 | -0,09 |
| uncultured Techofilosea | | -0,24 | -0,16 |
| Telaepolella | | -0,16 | -0,16 |
| D3P05A02 | | -0,09 | -0,24 |
| Sorosphaerula | | 0,05 | 0,13 |
| Filamoeba | | 0,05 | -0,16 |
| uncultured_eukaryote | | 0,05 | 0,13 |
| Mortierella | | 0,24 | 0,31 |
| uncultured Agaricales | | 0,35 | 0,35 |
| uncultured Cercomonadidae | | 0,38 | 0,31 |
| LKM11 | | 0,56* | 0,64** |
| Hygrocybe | | 0,67** | 0,67** |

Table S4: Kendall correlation coefficient for the enriched eukaryotic genera abundance regarding Burkholderia and Sphingobium abundances in ST soil. **p <* 0.05, ***p <* 0.01, ****p <* 0.001.

|  | Correlation with Burkolderia abundance | | Correlation with Sphingobium abundace |
| --- | --- | --- | --- |
| D3P05A02 | | -0,43 | -0,64* |
| Spumella | | -0,36 | -0,29 |
| Lobochlamys | | -0,29 | -0,21 |
| Colpoda | | -0,07 | -0,29 |
| Opisthonecta | | 0,24 | 0,24 |
| Chlamydomyxa | | 0,43 | 0,21 |
| Filamoeba | | 0,79 | 0,57 |
| D3P05A02 | | -0,43 | -0,64 |

Table S5: Kendall correlation coefficient for the enriched eukaryotic genera abundance regarding Burkholderia and Sphingobium abundances in LT soil. **p <* 0.05, ***p <* 0.01, ****p <* 0.001.
